# First landscape of binding to chromosomes for a domesticated mariner transposase in the human genome: diversity of genomic targets of SETMAR isoforms in two colorectal cell lines

**DOI:** 10.1101/115030

**Authors:** Aymeric Antoine-Lorquin, Ahmed Arnaoty, Sassan Asgari, Martine Batailler, Linda Beauclair, Catherine Belleannée, Solenne Bire, Nicolas Buisine, Vincent Coustham, Alban Girault, Serge Guyetant, Thierry Lecomte, Benoît Piégu, Bruno Pitard, Isabelle Stévant, Yves Bigot

## Abstract

*Setmar* is a 3-exons gene coding a SET domain fused to a *Hsmar1* transposase. Its different transcripts theoretically encode 8 isoforms with SET moieties differently spliced. *In vitro,* the largest isoform binds specifically to *Hsmar1* DNA ends and with no specificity to DNA when it is associated with hPso4. In colon cell lines, we found they bind specifically to two chromosomal targets depending probably on the isoform, *Hsmar1* ends and sites with no conserved motifs. We also discovered that the isoforms profile was different between cell lines and patient tissues, suggesting the isoforms encoded by this gene in healthy cells and their functions are currently not investigated.

## INTRODUCTION

*Setmar* is one of the 52 neogenes identified in the human genome that originate from a DNA transposon [Supplementary Table S1], i.e. from the exaptation of an open reading frame (ORF) encoding a transposase, the enzyme is able to carry out all the DNA cleavage and strand transfer steps required for the transposition of this kind of transposable element (TE) [1]. *Setmar* appeared about 40-58 million years ago in the anthropoid lineage at the origin of the hominoids and the old and new world monkeys [2]. The gene is located on the human chromosome 3 (positions 4,292,212 to 4,328,658 in GRCh38/hg38) and arose from two neighboring genes, the first comprising two exons coding a lysine methyltransferase (SET) and the downstream second coding a *Hsmar1* transposase (HSMAR1) belonging to the *mariner* family. *Mariners* are very simple TEs composed of one transposase ORF flanked by two inverted terminal repeats (ITRs). These flanking sequences are specifically bound by the transposase, that excises the TE from a 'donor' site before catalysing its insertion at another ('target') locus. During evolution of the anthropoid lineage, the accumulation of at least three mutations modified the expression properties of the SET and HSMAR1 genes. They acquired the ability to be transcribed in a single RNA transcript in which the SET domain can be N-terminally fused to the *Hsmar1* moiety after intron excision to produce the SETMAR protein (also called Metnase) [2].

Recent work on *Setmar* RNA transcript variants isolated from the hematologic neoplasms of patients [3] and mRNA sequences available in databases revealed a complex situation. Alternative transcripts have the potential to code for two SET isoforms (V4 and X4) and eight SETMAR isoforms ranging from 40 to 78 kDa, although only the largest SETMAR isoform contains a complete SET moiety (Figure 1, Supplementary Table S2 and data S1). The eight SETMAR transcripts originate from alternative transcription start sites located upstream, within or downstream of the region encoding the SET domain, together with alternative splicing in the second exon of the SET moiety. Other variants resulting from single nucleotide polymorphisms can be found in human populations ([3] and http://databases.lovd.nl/whole_genome/view/SETMAR). The detection of SETMAR in various colorectal [4], melanoma, breast cancer (Supplementary Figure S1), and leukaemia cell lines [5] confirmed that only five transcript variants were translated into protein isoforms (Figure 1, V1, V2, X2, V5 and an isoform with a molecular weight (MW) similar to that of HSMAR1), although differences exists from one cell line to another [4]. To our knowledge, it is not established which protein isoforms are present in healthy and cancerous tissues of patients.

**Figure 1.**
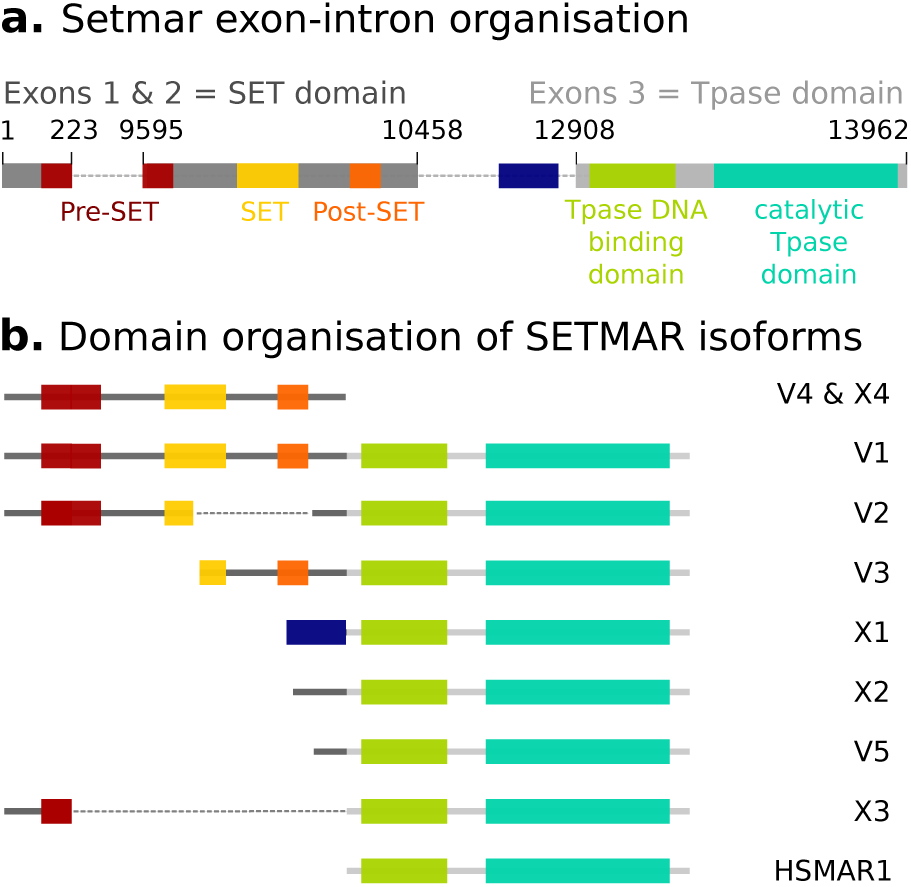
Organisation of the *Setmar* gene (**a**) and the various isoforms resulting from its transcription (**b**).

Since a decade, significant efforts have been carried out to elucidate the enzymatic activities of the SET and HSMAR1 moieties, to identify cellular protein partners, and to integrate this knowledge in biological pathways. In summary, the SET moiety was associated with the dimethylation of histone 3 lysines 4 and 36 [6] and the methylation of lysine 130 of the splicing factor snRNP70 [7]. The N-terminal domain of the HSMAR1 moiety has kept its ability to bind specifically to the ITRs of the *Hsmar1* transposon *in vitro.* Its C-terminal domain displays most of the cleavage and strand transfer activities of *mariner* transposases, except that the 3' second strand cleavage at the outer ITR ends is severely affected by a mutation of the catalytic triad (DDN instead of DDD) [8,9]. SETMAR has an additional DNA binding activity, in the form of a stable complex with the hPso4 protein [10,11], that is able to bind *in vitro* to non-ITR double-stranded DNA target. To date, it is not clear whether the different SETMAR isoforms can bind to chromosomal DNA in human cells and whether they target specific *Hsmar1* ITR related sequences and-or non-ITRs. From a functional standpoint, the biological activities of the SETMAR V1 domains (Figure 1) have been documented in the context of at least three housekeeping mechanisms, where the protein enhances chromosomal decatenation [12,13], improves the efficiency and the accuracy of DNA repair by non-homologous end-joining [14–17], and positively impacts the restart of stalled replication forks after DNA damage repair [18]. Interestingly, SETMAR V1 is also able to enhance the chromosomal integration of transfected DNA fragments [19] and of lentiviral DNA into host cells genome [20]. However, the precise molecular mechanism is unknown and whether the choice of integration sites is governed by the binding properties of SETMAR is still an open question. Also, it is not clear whether (and how) this mechanism is connected to metastatic processes, by mediating integration of the free DNA released by tumor cells into the chromosomal DNA of healthy cells [21,22].

Here, we derived the DNA binding landscape of several SETMAR isoforms in the colorectal cell lines SW48 and HT29. These lines display specific expression profiles of SETMAR isoforms: HT29 cells express a single SETMAR isoform (V2) whereas SW48 cells express several (V1, V2, X2, V5 and HSMAR1) [4]. We also revisit the annotation of the human genome for the *Hsmar1* element and its associated 80-pb miniature element, MADE1, using logol [23] and BLAST+, and we could detect and annotate ~2500 novel MADE1 copies. We found that SETMAR binding sites occur mostly at MADE1 sequences, although a significant fraction of them (up to ~50%) can also be found at unannotated regions. In addition, we document the expression profile of SETMAR isoforms in healthy and cancerous colorectal tissues. Strikingly, X2 and V2 are the only SETMAR isoforms expressed in both healthy and cancerous colon biopsies. We discuss which SETMAR isoforms are found in healthy tissues and tumors, with respect to the interest to study this protein in cancerous lines.

## MATERIALS AND METHODS

### Culture of cell lines

HeLa cells, human colorectal cancer cell lineages (SW48, SW403, HT29), human melanoma cell lineages (BRIS, SK-MEL28and 518-A2), and human breast cancer cell lineages (MCF7, MDA-MB231, SKBR3 and T47D) were all cultured in Dulbecco's modified Eagle's medium (DMEM) supplemented with 10 % fetal bovine serum (FBS), at 37 °C and with 5 % CO_2_. HeLa cell transfection with 1 μg of pVAX-HSMAR1 DNA was monitored as described [4].

### Samples of non-tumoral and tumoral colon tissues

Two samples of colon tissues, tumoral and adjacent non-tumoral tissues, were recovered from patients after surgery for a colorectal cancer in 2007 or 2008. Samples were stored at −80°C by the tumor bank of the Tours CHRU. Patients were informed of the possibility of the use of levies for research and their agreement to participate in this research was collected. Selected samples did not display more than 50% of tumoral cells (Supplementary Table S3).

### Protein extraction from non-tumoral and tumoral colon tissues

Tissue slices (10 μm) were generated with a cryomicrotome on frost samples of tumoral and non-tumoral tissues, and stored at −80°C. For each patient, protein lysates were made by suspending a tenth of slices in iced RIPA buffer (20 mM Tris-HCl pH7.2, 150 mM NaCl, 1 mM EDTA, 1% Glycerol, 1% Triton X100, 0.5% doxycholate, 0.1% SDS, 1X-Complete Protease Inhibitor Cocktail (Roche Applied Sciences, Meylan, France)), vortexed for 1 min, incubated on ice for 15 min, centrifuged at 15,000 g for 15 min at 4°C. After recovery of supernatants, proteins were quantified with a Quick Start™ Bradford Protein Assay (Bio-Rad, Richmond, USA) and conserved at −20°C.

### Antibody purification

Murine pre-immune and anti-HSMAR1 polyclonal sera were produced by DNA vaccination using ICANtibodies™ technology (In Cell Art, Nantes, France) as described [4]. Polyclonal antibodies (pA) contained in pre-immune and anti-HSMAR1 murine sera were purified using Protein A/G MagBeads (GenScript, Piscataway, USA). Their quality was then verified by polyacrylamide gel electrophoresis (PAGE) after staining with Coomassie blue and their concentration was defined using the BCA Protein quantification Kit (Interchim, Montluçon, France).

### PAGE, immunoblotting, and hybridization of antisera

Protein extracts from cultured cells, PAGE, transfer onto a polyvinylidene difluoride (PVDF) membrane, antibodies incubations and imaging with a FUJI LAS4000 imager were carried out as described [4].

### ChIP experiments

Chromatin samples were prepared from non-synchronous and exponentially growing SW48 and HT29 cells. Chromatin shearing was performed with a Bioruptor ultrasonicator (Diagenode, Belgium). Chromatin immunoprecipitation was performed with 10 μg of purified pre-immune or HSMAR1 pA, and purification of immunoprecipitated DNA was done using the iDeal ChIP-Seq kit following supplier's recommendations (Diagenode, Ougrée, Belgium). Libraries for Illumina sequencing were made using iDeal ChIP-Seq & Library Preparation Kit (Diagenode, Ougrée, Belgium). DNA quantities were monitored at various steps of the procedure with the Qubit® dsDNA HS Assay Kit (Molecular probes, Eugene, USA). Fragment size selection, library quality control and Illumina sequencing (HiSeq 51 nucleotides, TruSeq SBS Kit v3 (Illumina, Fulbourn, United Kingdom)) were achieved at the Imagif sequencing platform (CNRS, Gif-sur-Yvette, France), following published quality recommendations [24]. Data published in this paper are based on three biological replicates.

### ChIP-Seq Analyses

ChIP-Seq sequence reads were mapped to the human genome assembly hg38 (December 2013; available at http://www.ncbi.nlm.nih.gov/assembly/GCF_000001405.26) with the Bowtie short read aligner [25]. Peak calling was done with the peak-calling prioritization pipeline (PePr1.1.5) [26] from bam files, and normalized over input (three biological replicates).

### Data analyses

Annotation of *Hsmar1* and MADE1 copies in hg38 is described in Supplementary data 2. Unique or intersecting annotations were computed using bedtools. Ontologies and search for conserved motifs were carried out with GLAM2 [27], the GREAT pipeline [28], and the RSAT pipeline [29,30]. Data analyses and most graphic representation were done with R (https://www.r-project.org/). Graphic representations of chromosomes in the hg38 genome model were performed using svg files calculated with the DensityMap software at http://chicken-repeats.inra.fr/launchDM_form.php and gff files parameterized as recommended [31] in order to supply the chromosomes sizes, the locations of centromers, and a suitable description of the third field, i.e. the features in column 3. The locations of each centromer in chromosomes of the hg19 model were recovered and updated with liftover at the UCSC web site.

### Data repository

All raw and processed data are available through the European Nucleotide Archive under accession number PRJEB19196. The gff files describing the updated annotation of Hsmar1 and MADE1 copies in the hg38 release and the ChIP-Seq peaks were supplied as Supplementary data 3 and 4.

## RESULTS

### SETMAR binding sites along human chromosomes

Chromosomal targets of SETMAR were identified by ChIP-Seq on cell lines HT29 and SW48, using a custom made polyclonal antibody directed against the HSMAR1 domain of SETMAR. The specificity of this key antibody has been addressed previously which showed a good specificity/sensitivity ratio [4]. Our experience is that none of the commercial monoclonal and polyclonal antibodies commercially available and directed against the HSMAR1 domain are efficient are for ChIP experiments. ChIP-Seq of pan-HSMAR1/SETMAR was preferred to transient transfection of plasmids expressing individual SETMAR isoforms because transfected cell populations typically display heterogeneous expression levels that can span three orders of magnitude. This may not reflect the expression levels found *in vivo* [33] and can have strong impact on the binding landscape. Indeed, HSMAR1 and SET were found to accumulate at nucleoli (or periheterochromatin) when overexpressed in HeLa [33] and other cell lines (http://www.proteinatlas.org/ENSG00000170364-SETMAR/subcellular). Overall, we found 4366 and 8425 binding sites (BS) in HT29 and SW48 cells, respectively. Taking into account that in a few cases, peaks found in one cell type were split into two in the other, there is a total of 3513 peaks in common, together with 796 and 4908 only found in HT29 or SW48 cells.

Unfortunately, defining the co-occurrence of SETMAR BS with *Hsmar1* and MADE1 copies proved to be difficult, mainly because of their limited description in hg38 genome annotation. TEs annotation is notoriously difficult to carry out and it is not uncommon to assess its quality and re-annotate a particular family/subfamily. The quality of the RepeatMasker annotation was therefore verified and improved using logol and BLAST+ (Supplementary data 2). We produced a revised annotation describing 519 *Hsmar1* copies (displaying to 615 ITRs), together with 10295 MADE1 copies composed of 1875 full-length elements, 6246 elements truncated at one end (displaying a single ITR), and 2174 elements with damaged ITRs (outer ends lacking ≥ 5 nucleotides).

Intersections between ChIP-Seq peaks and the *Hsmar1* or MADE1 copies were calculated with bedtools using default parameters. Overall, there are 3549 (~83.9%) and 3998 (~49.6%) BS at *Hsmar1* or MADE1 copies in HT29 and SW48 cells, respectively, the vast majority of which (78.5%, 3449/4396) is common to both cell lines. Binding occurs mostly at MADE1 elements compared to *Hsmar1* ITRs, in both cell lines (Table 1). This result is consistent with the fact that SETMAR V2 is expressed in both cell lines (and likely other isoforms as well, in SW48 cells) and can bind to MADE1 and *Hsmar1* DNA *in vivo.* A total of 456 *Hsmar1* and 6062 MADE1 copies are not bound by SETMAR. We thus asked whether these copies were associated with lamina (LAD) or nucleolus (NAD) associated domains ([34] and http://biorxiv.org/content/early/2016/05/24/054908), but we could find no statistical correlation. Similarly, we could not find any association with intergenic regions, genes body, 5' proximal and 3' distal region of genes and introns.

**Table 1a.** Number and location of ChIP-Seq peaks in two colorectal cell lines

| Binding sites | HT29 cells | SW48 cells | Common in both cell lines |
| --- | --- | --- | --- |
| *Hsmar1* | 19 | 21 | 123 |
| MADE1 | 197 | 710 | 3326 |
| Other | 538 | 4082 | 163 |

**Table 1b.** Proportion of *Hsmar1* and MADE1 ITRs bound by SETMAR in two colorectal cell lines

|  | HT29 | SW48 |
| --- | --- | --- |
| Proportion of *Hsmar1* ITRs bound in hg38 | 23.2 % | 23.6 % |
| Proportion of MADE1 ITRs bound in hg38 | 34.2 % | 39.7 % |

We next characterized the BS common to both cell lines and overlapping with truncated MADE1 copies. Since they are composed of a single ITR, they minimize the confounding effect of nearby peaks. BS are centered on a sequence corresponding to positions 5 to 26 of *Hsmar1* and MADE1 ITRs (Figure 2a, Supplementary data 2). We note, however, that while residues located at position 8 to 17 and 21 to 26 are critical for HSMAR1 binding *in vitro* [2, 35], the conserved CG dinucleotide located at positions 24 and 25 appears less common at the genomic DNA level. On average, BS display 92.8 ±3.8% identical residues with the MADE1 consensus. In contrast, we found more sequence diversity in unbound MADE1 sequences, with an average of 87.8 ±5.2% identical residues.

**Figure 2.**
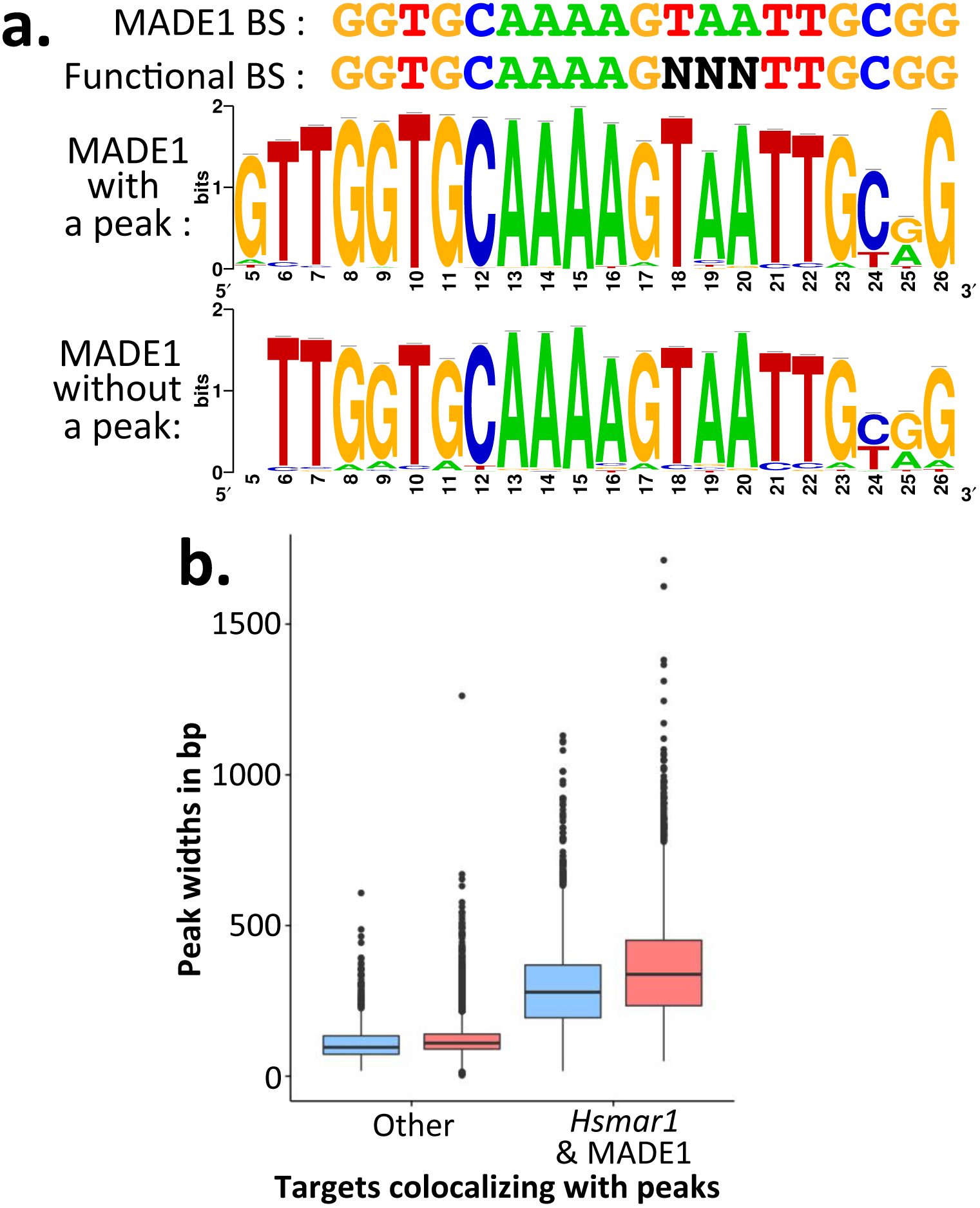
Graphic analyses of ChIP-Seq peak features. (**a**) Conserved motifs located by GLAM2 in truncated MADE1 copies overlapping or not with a ChIP-Seq peak. On the top, the SETMAR BS sequence in the MADE1 consensus sequence is shown, and below the sequence of the BS defined *in vitro* [2,35]. Positions along the ITR are indicated below the horizontal axes of Weblogos. The population of MADE1 without peaks was heterogeneous BS and since it is a mixture of elements with few conserved BS and elements with an ability to be bound by SETMAR isoforms that depends on the local chromatin structure. (**b**) Boxplots representing the distributions of the width (in bp) of peaks colocalizing with *Hsmar1* or MADE1 ITRs (2 boxes on the right) or elsewhere in the human genome (2 boxes on the left) depending on the cell lines (HT29 in blue, SW48 in red).

Thus, our results support well the fact that SETMAR V2 (and likely of other isoforms) binding to MADE1 and *Hsmar1* DNA depends on the sequence conservation of their ITR, but it also depends on cell specific host factor(s).

### Features of non-ITR binding targets along human chromosomes

The number of BS outside of *Hsmar1* or MADE1 copies was higher in SW48 than in HT29 cells, which correlated with the presence of four other SETMAR isoforms in SW48 (V1, X2, V5 and HSMAR1) that were dramatically more abundant than or absent in HT29 cells. On average, these BS are associated to lower q-values and smaller widths than those found at *Hsmar1* or MADE1 DNA BS in HT29 and SW48 cells (Supplementary Figure S2; Figure 2b). Altogether, this suggests that they require a specific molecular machinery, which may be independent of the presence of hPSO4 in both cell lines, as this protein is present in similar amounts in both cell lines (Figure 3).

**Figure 3.**
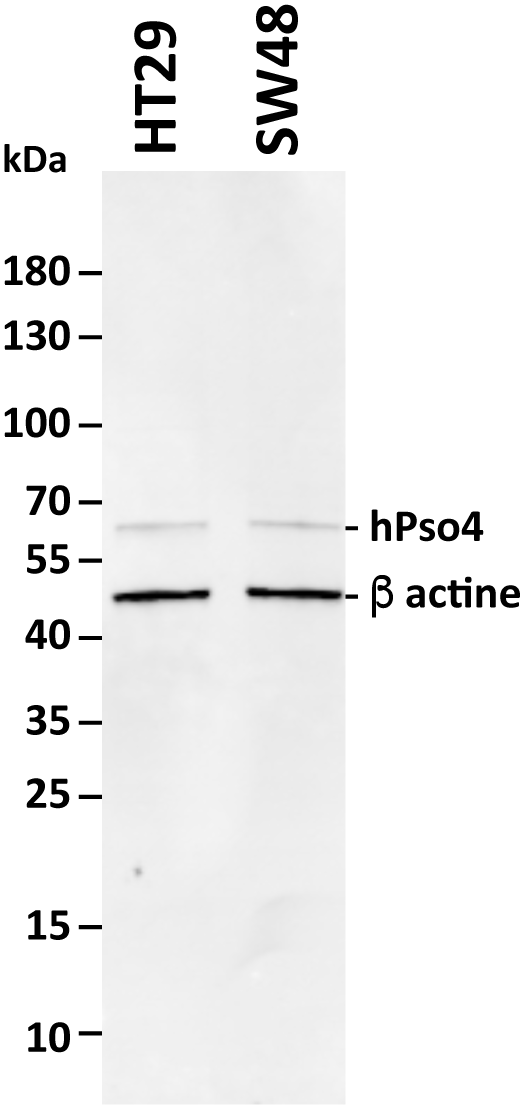
Control of the hPso4 (also known as PRPF19) presence in HT29 and SW48 cell lines. Protein extracts (50 μg) were used to make Western blot. hPso4 and beta actin detections were achieved together using a rabbit anti-human hPso4 (PA5-24797, ThermoFischer Scientific) and a rabbit anti-human beta actin (EPR16769, Abcam) as primary antibodies and donkey anti-rabbit IgG-IR DYE800LT as a secondary antibody. Incubation with both antibodies were done overnight at 4°C in the Odyssey blocking buffer TBS (LI-COR). Three washings (10 min) of membranes were done in PBS1X, 0.1% triton X100, at room temperature after each antibody incubation. Membranes were analysed using an Odyssey CLx Imaging system (LI-COR). Protein molecular weights are indicated in the left margin.

Searches for common ontologies and conserved transcription factor (TF) BS were conducted using the GREAT pipeline with default parameters (Supplementary data 5). We could not find significant association between BS at *Hsmar1/*MADE1 sequences, gene function or their location with respect to genes. For BS located outside of Hsmar1/MADE1 sequences, 200 of the 4783 these BS could be associated to genes involved in epithelia and mesenchymal stem cell biology.

As expected in term of genomic distribution, BS localized at ITRs followed that of *Hsmar1/*MADE1 copies. Unexpectedly, however, more than 75% of the BS located outside of *Hsmar1*/MADE1 sequences were not distributed at random and were clustered in a limited number of regions, ranging from 1 to 20 Mbp, found in only 10 chromosomes (Figure 4). Although no common DNA motif could be found, BS characteristics and their genomic distribution along chromosomes certainly reflect the (direct or indirect) specific binding of some SETMAR isoforms to DNA or chromatin. We next searched for putative transcription factor binding motifs (TFBM) at BS located outside of Hsmar1/MADE1, with the RSAT pipeline (Supplementary data 5). We found a clear signature of EGR1 and-or MEF2 TFBM in 69% of them, suggesting possible physical interactions between the two TFs and SETMAR isoforms.

**Figure 4.**
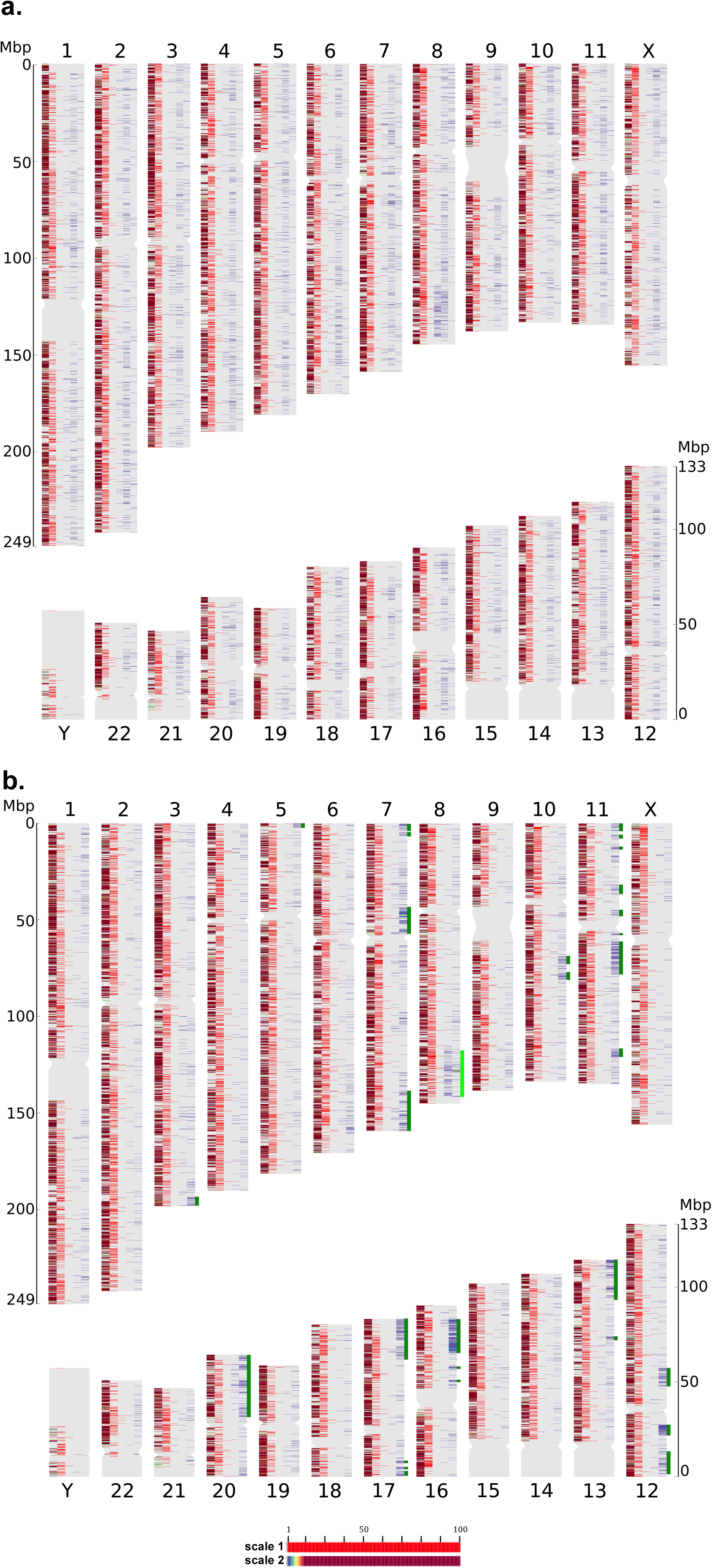
Distribution of gene densities, *Hsmar1* and MADE1 copies and peaks in chromosomes of the hg19 genome model. (**a**) Graphic representation of the gene density, and the occurrences of MADE1 and *Hsmar1* copies and ChIP-Seq peaks colocalized with a *Hsmar1* or a MADE1 copy along the human chromosomes. Each chromosome is represented from the left to the right in six columns: the gene density, the occurrences of MADE1 copies, the occurrences of *Hsmar1* copies, the occurrences of ChIP-Seq peaks co-localized with a *Hsmar1* or a MADE1 copy only in HT29 cells, both in HT29 and SW48 cells, and only in SW48 cells. (**b**) Graphic representation of the gene density, and the occurrences of MADE1 and *Hsmar1* copies and the ChIP-Seq peaks that were not co-localized with a *Hsmar1* or a MADE1 copy along the human chromosomes. Each chromosome is represented from the left to the right in five columns: the gene density, the occurrences of MADE1 copies, the occurrences of *Hsmar1* copies, the occurrences of ChIP-Seq peaks that did not co-localize with a *Hsmar1* or a MADE1 copy only in HT29 cells, and only in SW48 cells. The most concentrated region of non-ITR binding targets in HT29 and SW48 is indicated with a light green bar. In both graphics, gene density and ChIP-Seq peaks were colourized using the colour scale 1, the occurrences of MADE1 and *Hsmar1* copies used the colour scale 2. Densities and occurrences were calculated per window of 10^5^ bp. The centromer of each chromosome is located by a constriction in the shape of each chromosome.

### Expression of SETMAR isoforms in colon biopsies

Due to the heterogeneity of SETMAR isoform profiles between cancerous cell lines ([4,13] and Supplementary Figure S1) and that the expression profiles of healthy tissues or cancerous tumors can drift in primary cultures as well as in established cell lines [36–38], the presence of SETMAR isoforms was probed in biopsies of non-tumoral and tumoral colon tissues from 26 patients affected by a colon cancer. As a control, we also used a colorectal biopsy from a patient with no colorectal disorder. In healthy and tumoral colon tissues, we found that four (V1, V2, V5 and HSMAR1) of the five isoforms present in HT29 and SW48 cells were not detected but two other SETMAR isoforms, X2 and V3, were present (Figure 5a). The most abundant was the X2 isoform. Its expression level varies from 1 to 5-folds, depending on the nature of the biopsy, and the patient but not sex (Figure 5b). This isoform is also found in healthy tissues. The second protein, which was poorly expressed only in a few patients, has a molecular weight of 55 kDa that corresponds to the V3 isoform. Overall, the stark contrast between the SETMAR expression profile *in vivo* (healthy/cancerous tissues) and *in vitro* (established colorectal cell lines) shows that it is very sensitive to the cellular context. Interestingly, V2, X2 and V3 isoforms likely have altered or no SET catalytic activity since important parts of this domain are missing in these isoforms. Thus, the resulting proteins would be equivalent to a HSMAR1 tagged with N-terminal peptides that might modulate their interactions with putative (chromatin) partners.

**Figure 5.**
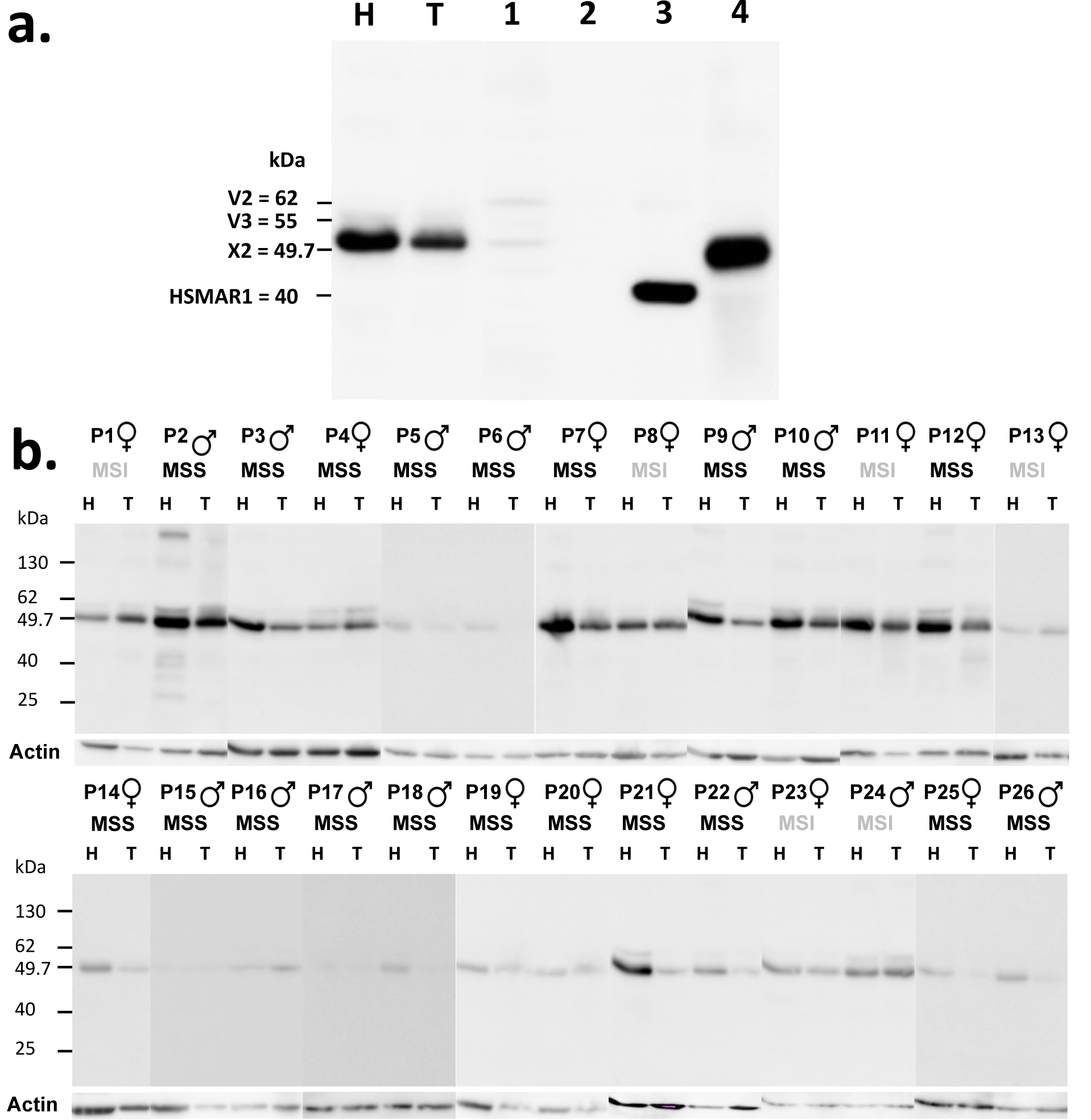
Western blot analyses of SETMAR isoforms in healthy and tumorous colorectal tissues of patients. (**a**) Protein extracts of two biopsies from patient 12 (P12; healthy (H) and tumoral (T) tissues, 50 μg), SW403 cell line that mainly expresses V2 and X2 SETMAR isoforms [4] (lane 1, 50μg), HeLa cells (lane 2, 50 μg), HeLa cells transfected with pVAX-Hsmar1 (lane 3, 50 μg), and a colorectal biopsy from a patient with no colorectal disorder (lane 4, 50 μg). Molecular weights of isoforms are indicated in the left margin. (**b**) Protein extracts from healthy (H) and tumoral (T) tissue biopsies of 26 patients (P1 to P26). Actin is shown as an internal loading reference. The sex of each patient and the phenotype of their colon cancer (microsatellite stable (MSS) and microsatellite instable (MSI)) are indicated. For imaging the Western blot results with the anti-Hsmar1 antibodies in (**a**) and (**b**), an exposure time of 1 hour was used while only 5 to 10 minutes were required for the actin controls (Abcam, ab13822). Protein molecular weights are indicated in the left margin.

## DISCUSSION

Our work shows for the first time that SETMAR binds at a variety of BS *in vivo.* As expected, binding occurs at the ITR of *Hsmar1* and MADE1, but also occurs at other sites centered on GC-rich regions with no obvious conserved motif. Remarkably, in HT29 cells, V2 binding occurs despite abundant hPSO4 levels in the nucleus. Overall, this novel population of BS unlikely result from the direct binding of SETMAR to chromosomal DNA, but may be mediated by other protein partners, possibly EGR1 and/or MEF2, that would interact through the pre-SET and/or the HSMAR1 domain.

The V2 isoform lacks intact SET and post-SET domains, which rules out indirect binding mediated by protein complexes involved in replication or DNA repair [3,5,18]. They would therefore result from interactions with DNA binding proteins that were so far not described and which would occur with the pre-SET subdomain and/or the HSMAR1 domain. In SW48, because the X2 and V5 isoforms and HSMAR1 display no SET subdomains, it is likely that these proteins have interactions with chromosomal DNA that are similar to those of V2. Under this hypothesis, most of the peaks unrelated to *Hsmar1* and MADE1 copies would therefore result from interactions between the V1 isoform and other protein partners, putatively EGR1 and MEF2. These peaks would likely not result from interactions between V1 and protein complexes involved in replication or DNA repair. Indeed, they are discrete and well defined, what is not expected in such interactions that are not specific in location in non-synchronized cell populations.

Our work, although limited to a cohort of 26 human samples, already provides important new data. Remarkably, our results demonstrate that the study of SETMAR functions using cell lines is tricky, especially for the V1 isoform that was so far never described as being present in healthy organs or in tumors *in vivo.* Here, our study was limited to 26 patients in whom only the V3 and X2 isoforms were detected in non-tumoral and certain cancerous biopsies. Currently, we have analysed 20 cell lines derived from colorectal cancers (13; [4] and Figure 2A, lane 1), melanoma (3) and breast cancers (4) (Supplementary Figure S1). In none of them the V3 isoform was detected while the V5 isoform was found in 7 colorectal lines. The V2 and V3 isoforms display both a damaged SET domain likely unable to bind to DNA. It is therefore reasonable to hypothesize that the ability of V3 to bind to chromosomal DNA is close to that of V2 and mainly occurred on ITR of *Hsmar1* and MADE1 copies. Currently, it cannot be discarded that most non-Hsmar1/MADE1 BS are due to the binding of V1, a SETMAR isoform that was so far only detected in a few cell lineages.

Since HSMAR1 3' end cleavage activity is severely impaired [8], HSMAR1 and isoforms like V2, V3, X2 and V5 are unlikely to mediate *Hsmar1/*MADE1 excision. However, HSMAR1 is still capable of mediating 5' end cleavage and DNA integration [8,19]. This raises the interesting possibility that SETMAR isoforms might catalyse integration of extracellular DNA, because circulating free DNA is often released from cell death or infection and can be passively transferred into cells [21,22]. Given the limited number of possible integration sites, mostly located outside of genes, SETMAR would then represent a nuclear mechanism protecting cells against the genotoxic effects of integration of circulating cell free DNA. This would represent a recent functional innovation, restricted to the anthropoid lineage.

## FUNDINGS

This work was funded by the C.N.R.S., the I.N.R.A., and the GDR CNRS 2157. It also received funding from a Research Program grant from the Cancéropôle Grand-Ouest, the Ligue Nationale Contre le Cancer and grants from Amgen and the French National Society of Gastroenterology.

